## Supplemental file for "Imaging Microglia Surveillance during Sleep-wake Cycles in Freely Behaving Mice"

### Supplementary information

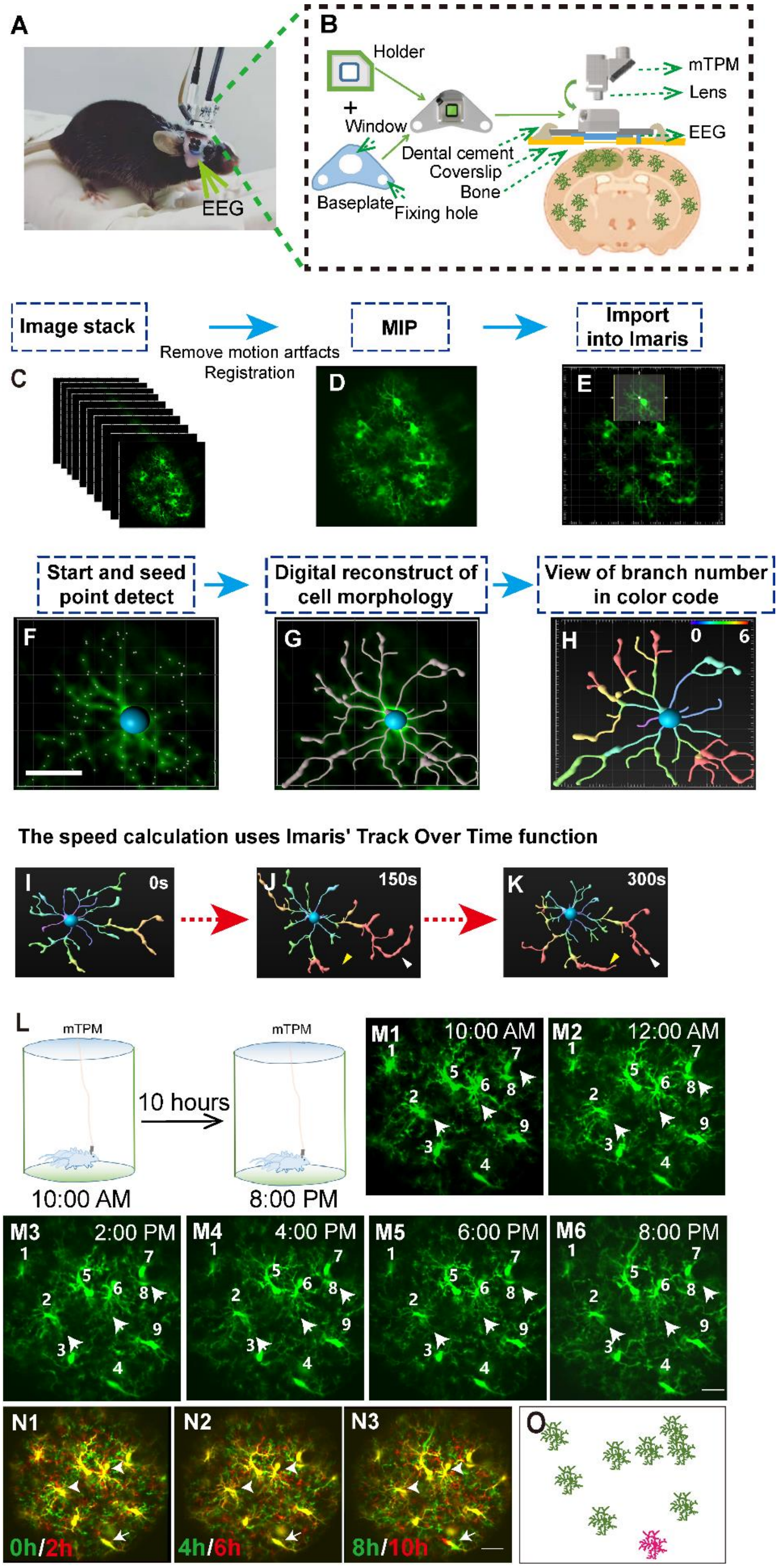

**Figure S1. Microglia dynamics imaging and analysis in freely behaving mice. Related to Figure 1.**

(A) Head-mounting of the mTPM. Picture of a free moving mouse with a head-mounted mTPM and EEG/EMG recording electrodes.

(B) Cartoon illustration showing that the mTPM headpiece was mounted onto an assembly consisting of a baseplate plus a holder glued onto a glass coverslip over a cranial window. Initially, the headpiece was screw-fastened to its holder, and their placement on the baseplate was fine-adjusted using a triaxial motorized stage when the animal was immobilized on an imaging stage. The holder was then fixed to the baseplate with dental cement once a proper FOV was located. The animal could then be released and left to move freely in a mouse cage. After an imaging session, dismounting of the headpiece was made easy by unscrewing and unplugging it from the holder.

(C-K) Pipeline for microglial surveillance analysis.

(C-D) An image stack (5-min, 1500 frames) pre-processed using ImageJ software to correct for motion artifacts and to align them using the StackReg plugin. In the case of multi-plane imaging, consecutive z-stacks were first each compressed to single xy projections. A maximum-intensity projection (MIP) was created from the final stack for morphological analysis.

(E) Importing the MIP image to Imaris software and setting ROIs (region of interest) to encompass selected cells with clear cell bodies and processes.

(F-H) For a given cell, Autopath's algorithm was used to find the right starting point and seed point. The radius of process was measured by the shortest distance algorithm. Microglial processes were digitally reconstructed from the calculated seed point and process radius. Each branch was color-coded by its branch point number.

(I-K) Process end-point speed was calculated through Imaris' track-over-time function, the average absolute speed is given by track length divided by time of tracking. Yellow and white arrowheads in (G) and (K) show elongating and contracting process ends, respectively. Scale bar, 20 μm.

(L-O) An example illustrating long-term imaging of microglia in freely behaving mice.

(L) Experimental setup. While the animal was roaming freely in the cage, time-lapse mTPM images were obtained continually at 5 FPS, starting from 10:00 AM and ending at 8:00 PM.

(M) mTPM images of microglia at different time points. Note that no signs of photodamage or photobleaching caused by long-enduring *in vivo* mTPM imaging. All images shown correspond to the wake states of the animal. Numerals identify individual microglia.

(N-O**)** Overlay of images in (M) paired at 2h time intervals. Arrows mark the translocation of a cell body (cell number 4). Cartoon in (O) shows that only 1 out 9 microglia cells exhibited local motility (pink) with the rest (green) position unchanged over the duration of observations. Scale bar, 30 μm.

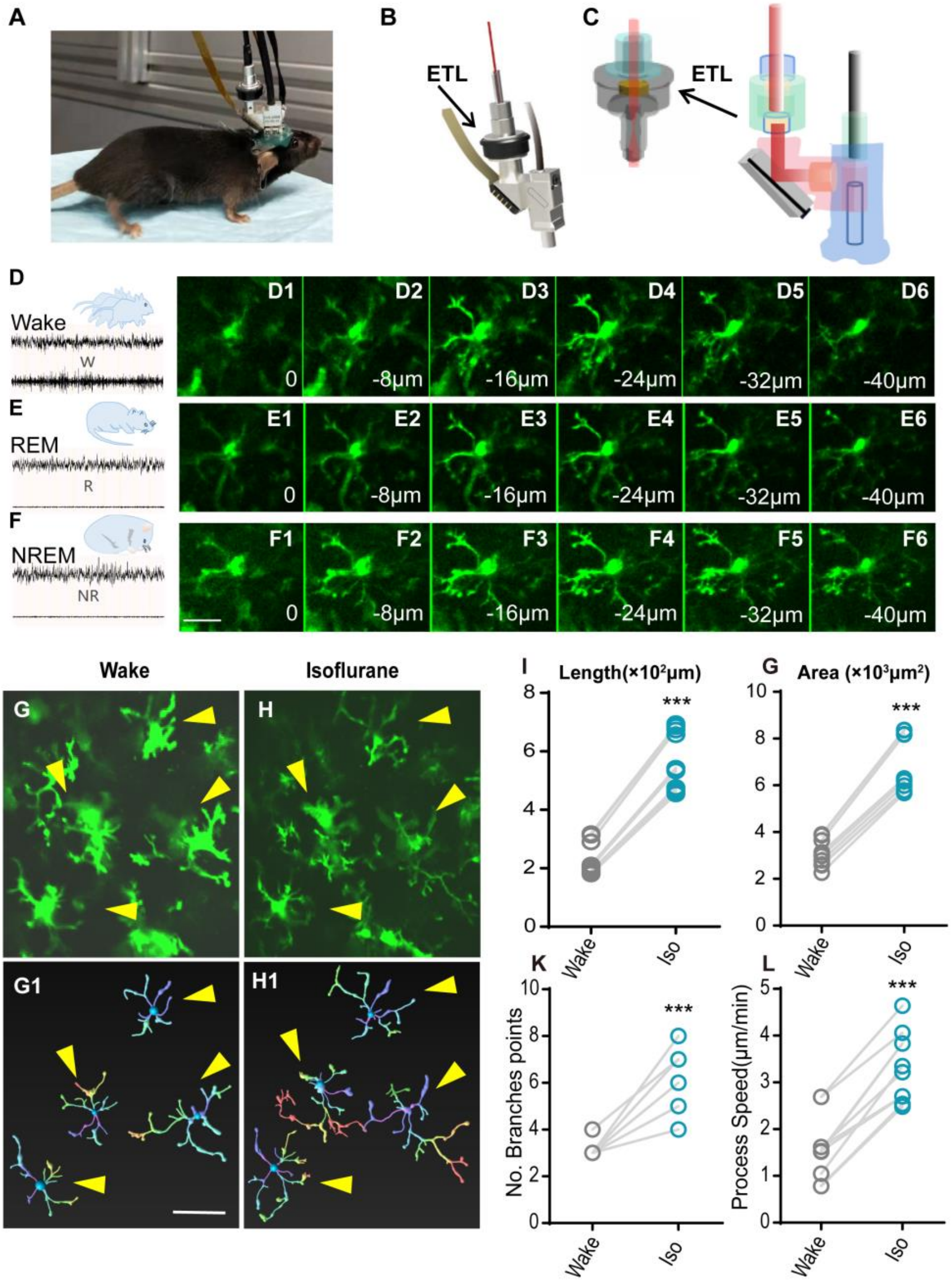


**Figure S2. Multiplane imaging of microglial surveillance and changes of microglial surveillance under anesthesia. Related to Figure 2.**

(A) Freely behaving mouse wearing an mTPM integrated with an electrical tunable lens (ETL).

(B-C) Schematic of the ETL assembled into the headpiece. The ETL (EL-3-10, Optotune, Dietikon, Switzerland) provided a Z-level change of ~45 μm, allowing for multi-plane imaging.

(D-F) Multi-plane imaging at different wake-sleep states: wake (D), REM (E), and NREM (F). The z-stack could then be used for correction of z-drift artifacts and for 3D analysis of microglial surveillance. Scale bar, 15 μm.

(G-L) Changes of microglial surveillance under anesthesia.

(G-H) Representative micrographs of microglia under wake and anesthetized states. (G1-H1) Digitally reconstructed microglial morphology corresponding to G-H. (I-L) Changes of microglial process length (I), area (G), number of branch points (K), and process end-points speed (L) induced by anesthesia. paired *t*-test. Scale bar, 30 μm. ***$P < 0.001$, Wake versus Iso. $n = 12$ cells from 3 mice for each group.

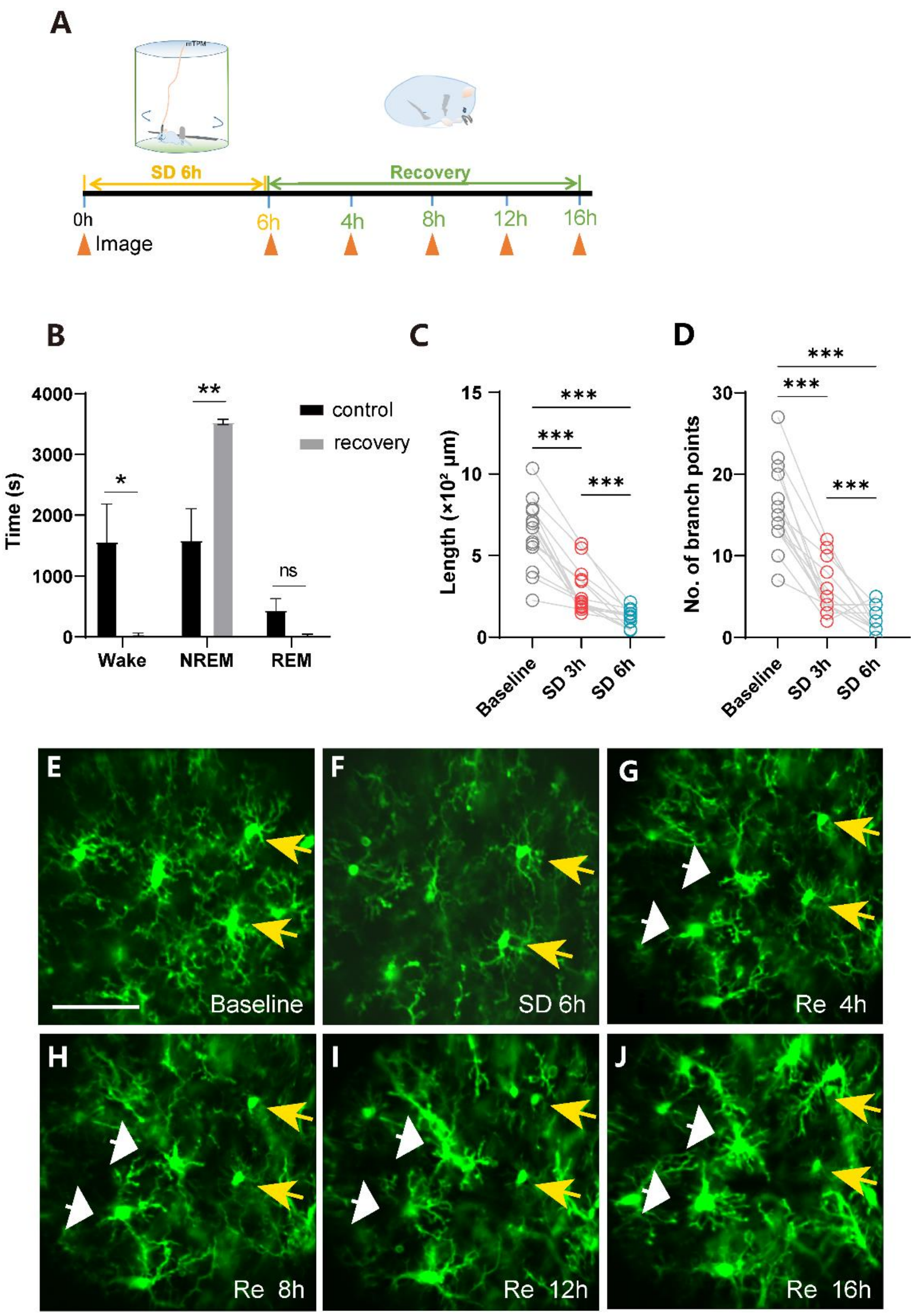


**Figure S3. Changes of microglial surveillance in the state of sleep deprivation and recovery. Related to Figure 3.**

(A) Experimental setup for sleep deprivation and recovery.

(B) Altered sleep-wake homeostasis during recovery from sleep deprivation (zeitgeber time 2-8). Recovery was during zeitgeber time 9-10. unpaired *t*-test. ns, not significant, $^{*}P < 0.05$, $^{**}P < 0.01$. $n = 4$ mice for each group.

(C-D) Quantitative analysis of microglial length (C) and number of branch points (D) based on multi-plane microglial imaging. One-way ANOVA with Tukey's post-hoc test. $n = 14$ cells from 3 mice for each group.

(E-J) Microglial processes contracted after sleep deprivation (SD) and partially restored during recovery (Re), baseline (E), SD 6h (F), Re 4h (G), Re 8h (H), Re 12h (I), and Re 16h (J). Scale bars, 30 μm.

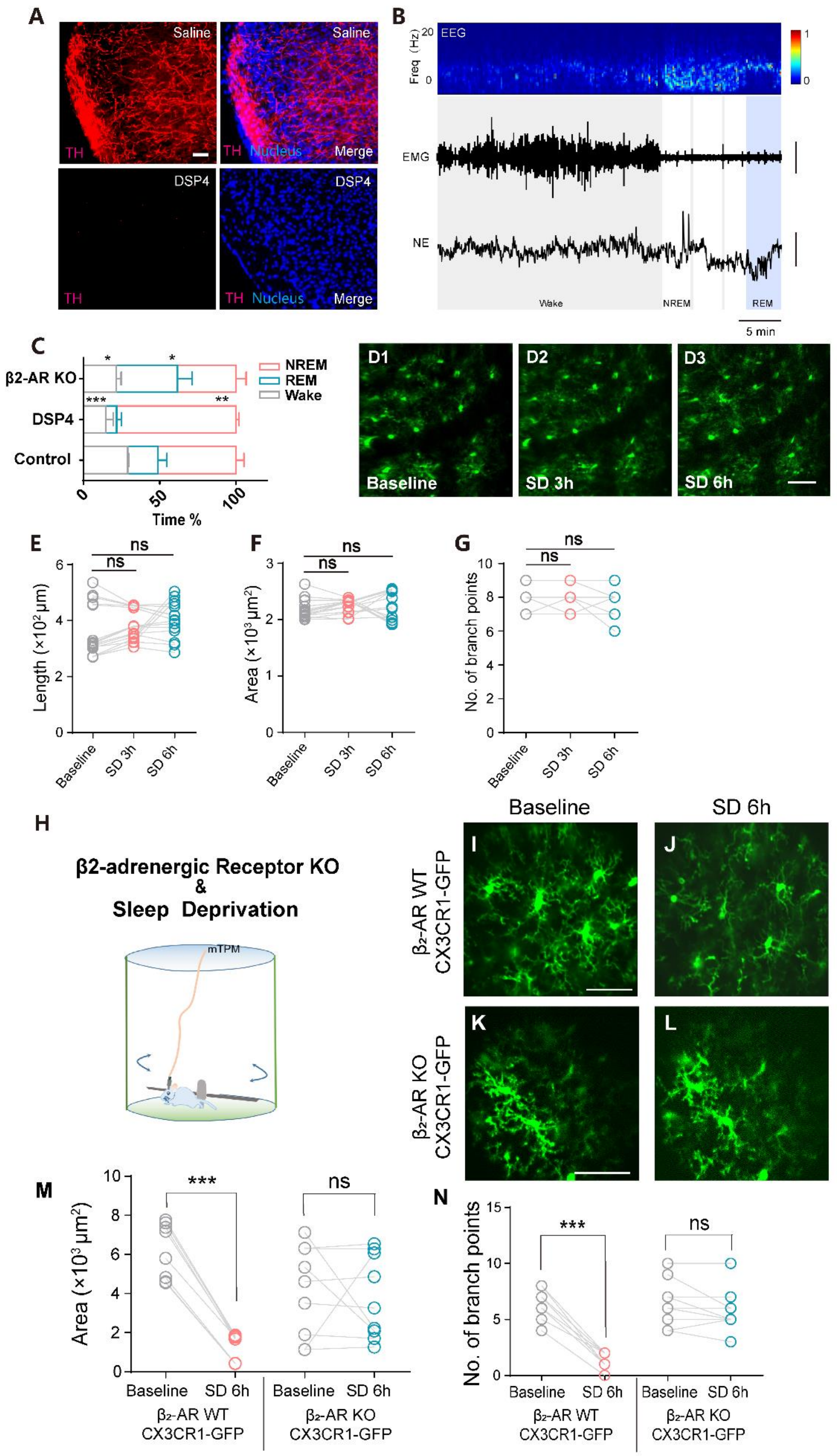
A
Saline
Saline
TH
TH Nucleus Merge
DSP4
DSP4
TH
TH Nucleus Merge
B
EEG
Freq (Hz)
20
0
1
0
EMG
NE
Wake
NREM
REM
5 min
C
β2-AR KO
DSP4
Control
NREM
REM
Wake
Time %
0
50
100
D1
D2
D3
Baseline
SD 3h
SD 6h
E
Length (×10² μm)
ns
ns
Baseline
SD 3h
SD 6h
F
Area (×10³ μm²)
ns
ns
G
No. of branch points
ns
ns
H
β2-adrenergic Receptor KO
&
Sleep Deprivation
mTPM
Baseline
SD 6h
I
J
K
L
β2-AR WT CX3CR1-GFP
β2-AR KO CX3CR1-GFP
M
Area (×10³ μm²)
***
ns
Baseline
SD 6h
Baseline
SD 6h
β2-AR WT CX3CR1-GFP
β2-AR KO CX3CR1-GFP
N
No. of branch points
***
ns
Baseline
SD 6h
Baseline
SD 6h
β2-AR WT CX3CR1-GFP
β2-AR KO CX3CR1-GFP

**Figure S4 Altered sleep-wake states after DSP4 administration and β$_2$AR knockout and controlling microglial surveillance during SD by LC-NE signal. Related to Figure 5.**

(A) Inhibition of tyrosine hydroxylase (TH) expression in somatosensory cortex of mice treated with DSP4. Nuclei were stained with DAPI, and saline was used in the control group. Scale bars, 50 μm.

(B) Representative traces of simultaneous multimodal recordings in the somatosensory cortex during the sleep-wake cycle in freely behaving mice treated with DSP4. From top to bottom: EEG and its power spectrogram (0-20 Hz); EMG (scale, 0.2 mV); NE signals reflected by the z-score of the $GRAB_{NE2m}$ fluorescence (scale, 2 z-score). The brain states are color-coded (wake, gray; NREM, white; REM, blue).

(C) DSP4 treatment and β$_2$AR knockout both reduced the wake time compared to controls. DSP4 significantly increased NREM time whereas β$_2$AR KO led to more REM time. *$P < 0.05$, **$P < 0.01$, ***$P < 0.001$ vs control, $n = 3$ mice for each group, unpaired *t*-test (two-tailed).

(D-G) Lack of microglial surveillance in LC-axon ablated animals. D: Representative mTPM images of microglia during SD. E-G: Statistics for microglial length (E), surveillance area (F) and number of branch points (G) in LC-axon ablated mice.

$n$ =15 cells from 3 mice for each group; ns, not significant, Friedman test with Dunn's post-hoc test.

(H-N) Lack of SD-dependent changes in microglial surveillance in the absence of $\beta_2$AR signaling.

(H) Experiment setup. (I-J) Representative mTPM micrographs from a (Cx3cr1-GFP+/-; Adrb2+/+) mouse. (K-L) Representative mTPM micrographs from a (Cx3cr1-GFP+/-; Adrb2-/-) mouse. No significant morphological changes were observed in $\beta_2$AR KO mouse subjected to SD. (M-N) Quantitative analysis of changes in microglial surveillance area (M) and number of branch points (N) in WT and $\beta_2$AR KO mice under baseline (wake state) and SD conditions. Scale bars, 30 μm.

**$P < 0.01$, wilcoxon test (two-tailed), Baseline versus SD 6h in WT mice in (**M**); ***$P < 0.01$, paired *t*-test (two-tailed) in (**N**). $n = 9$ cells from 3 mice for each group; ns, not significant
